# Automated detection of extracellular action potentials from single neurons

**DOI:** 10.1101/2022.06.06.494896

**Authors:** Zhuowei Cheng, Elmer Guzman, Tjitse van der Molen, Tal Sharf, Paul K. Hansma, Kenneth S Kosik, Linda Petzold, Kenneth R Tovar

## Abstract

Multi-electrode arrays (MEAs) non-invasively record extracellular action potentials (eAPs, also known as spikes) from hundreds of neurons simultaneously. However, because extracellular electrodes sample from the local electrical field, each electrode can detect eAPs from multiple nearby neurons. Interpreting spike trains at individual electrodes of high-density arrays requires spike sorting, a computational process which groups eAPs from single ’units’ based on assumptions of how spike waveforms correlate with different neuronal sources. Additionally, when experimental conditions result in changes to eAP waveforms, spike sorting routines may have difficulty correlating eAPs from multiple neurons at single electrodes before and after such waveform changes. We present here a novel, empirical method for unambiguously isolating eAPs from individual, uniquely identifiable neurons, based on automated multi-point detection of action potential propagation. This method is insensitive to changes in eAP waveform morphology because it makes no assumptions about the relationship between spike waveform and neuronal source. Our algorithm for automated detection of action potential propagation produces a ’fingerprint’ that uniquely identifies those spikes from each neuron. By unambiguously isolating eAPs from multiple neurons in each recording, on a range of platforms and experimental preparations, our method now enables high-content screening with contemporary MEAs. We outline the limitations and strengths of propagation-based isolation of eAPs from single neurons and propose how our automated method complements spike sorting and could be adapted to *in vivo* use.

## Introduction

Extracellular recording of action potentials has been used for decades, and has revealed fundamental insights in cellular and sensory neuroscience (Katz and Miledi, 1965; Hubel, 1957; Hubel and Weisel, 1963). The relative ease of use and the non-invasive nature of extracellular recording makes this technique technically approachable for recording action potentials from groups of excitable cells, compared to other electrophysiological recording configurations. Recent progress in fabrication and surface chemistry have led to the production of 2- and 3-dimensional extracellular electrode arrays, consisting of hundreds to thousands of densely-packed electrodes (Ju et al., 2015; Jun et al., 2017). Arrays such as these have been used to record simultaneously from hundreds of cultured primary neurons or iPS-derived neurons grown on these substrates (Wainger et al., 2014; Slomowitz et al., 2015), as well as from intact brains (Kodama et al., 2018; Berényi et al., 2014). However, the burden of increased data handling and computational cost accompany the increase in data channels, particularly when experiments require significant post-acquisition data processing (Einevoll et al., 2012: Hilgen et al., 2017).

Because extracellular electrodes sample from the electric field, rather than directly from each neuron, these electrodes detect eAPs from any proximal neuron. The likelihood that each electrode detects eAPs from multiple neurons increases with the density of neurons, compromising an electrode’s ability to isolate eAPs from single sources. Computational approaches collectively known as spike sorting have been developed to isolate, or sort, eAPs into discreet source ’units’ (Hill et al., 2011; Quiroga 2012). Multiple sorting routines have been introduced to automate data processing from arrays with several thousand electrodes (Hilgen et al., 2017; Diggelmann et al., 2018; Yger et al., 2018). Many such sorting routines assume that characteristics of the eAP waveforms from each electrode correlate with properties from each source neuron (Hill et al., 2011; Einevoll et al., 2012; Quiroga 2012; Harris et al., 2016). These assumptions are multiplied in recordings with thousands of electrodes, from neurons at high density. In different experimental preparations, ground truth validation of how sorting routines isolate spikes can only be achieved with technically challenging experiments that, for example, pair extracellular recording with intracellular or juxtacellular recording (Yger et al., 2018; Neto et al., 2016; Allen et al., 2018; Marques-Smith et al., 2018). Additionally, eAP waveforms from individual neurons can vary significantly within a single recording session due to changes, for example, in firing frequency (Buzsáki et al., 1996, Stratton et al., 2012). Common experimental manipulations that change eAP waveforms make it challenging for sorting routines to follow the same isolated ’unit’ between experimental manipulations (Quirk and Wilson, 1999; Hilgen, 2019). Efforts to validate the assumptions intrinsic to spike sorting routines are ongoing (Magland et al., 2020).

However, the ability of any sorting routine to reliably isolate spikes from the same neuronal source during experiments that significantly change waveform shape remains largely untested.

We previously used detection of action potential propagation among extracellular electrodes to uniquely identify, isolate and characterize propagating eAPs from individual cultured neurons (Tovar et al., 2018). Because axonal action potential propagation is unidirectional, the sequence of eAP occurrence at each array electrode along the propagation path creates a ’fingerprint’ that identifies each detected neuron. Here we present the automation of this empirical approach to isolating eAPs from multiple uniquely identifiable neurons in large electrode arrays. Detection of eAP propagation among extracellular electrodes categorizes ensemble spiking data into eAPs from unique source neurons, and for the first time routinely provides ground truth for the isolation of eAPs in extracellular recordings. Because our algorithm is based on the eAP propagation path across multiple electrodes, it reliably detects the same source neuron between experimental conditions that affect spike morphology (Debanne et al., 2011), including high spiking frequency, temperature changes or drugs such as K^+^ channel blockers. Our method reveals multiple unique neurons in recordings from primary cultures grown on low- or high-density arrays, as well as from whole human brain organoids recorded with shank electrode arrays. Automated detection of propagation separates propagating eAPs in identified source neurons from all other spikes in any extracellular recording and enables tracking single neurons across experimental conditions. Our algorithm enhances and streamlines post-acquisition data handling because propagating eAPs from single neurons seen at multiple electrodes are redundant in terms of extracting a neuron’s spiking pattern. Our algorithm can also be used in combination with traditional spike sorting routines and redundant eAPs identified by our algorithm can be ignored by spike sorting routines. Details of use of our algorithm, along with documentation, can be found at https://github.com/ZhuoweiCheng/Propagation-Signal-and-Synaptic-Coupling-Algorithm.

## Results

### Detection of action potential propagation identifies single neuron eAPs

In this work we used previously characterized features of action potential propagation detection on MEAs (Tovar et al., 2018) to design an algorithm that automates isolation of spikes from single neurons. Action potential propagation is evident in arrays of extracellular electrodes by the repeated co-occurrence of eAPs at multiple electrodes (Bridges et al., 2017; Tovar et al., 2018). The propagation sequence among cohort electrodes identifies such eAPs as originating from each unique source neuron.

Every recording from primary neurons cultured on arrays we examined had multiple unique examples of action potential propagation in multiple neurons, reflected by eAPs among electrode cohorts. Figure 1A shows a portion of an electrode array with black circles indicating a cohort of electrodes at which propagating action potentials from a single neuron were detected. The unidirectionality and high fidelity of action potential propagation results in the invariant sequence detection (Figure 1B) and low inter-electrode latency variability of these eAPs (Figure 1C). In this example, manual analysis showed that 93.3% (1939/2078) of the eAPs in electrode 1 were followed by eAPs in electrode 2, with a mean latency of 0.255 ± 0.018 ms. Similarly, 92.0% (1912/2078) of the eAPs at electrode 1 were followed by eAPs at electrode 3, with a mean latency of

**Figure 1.**
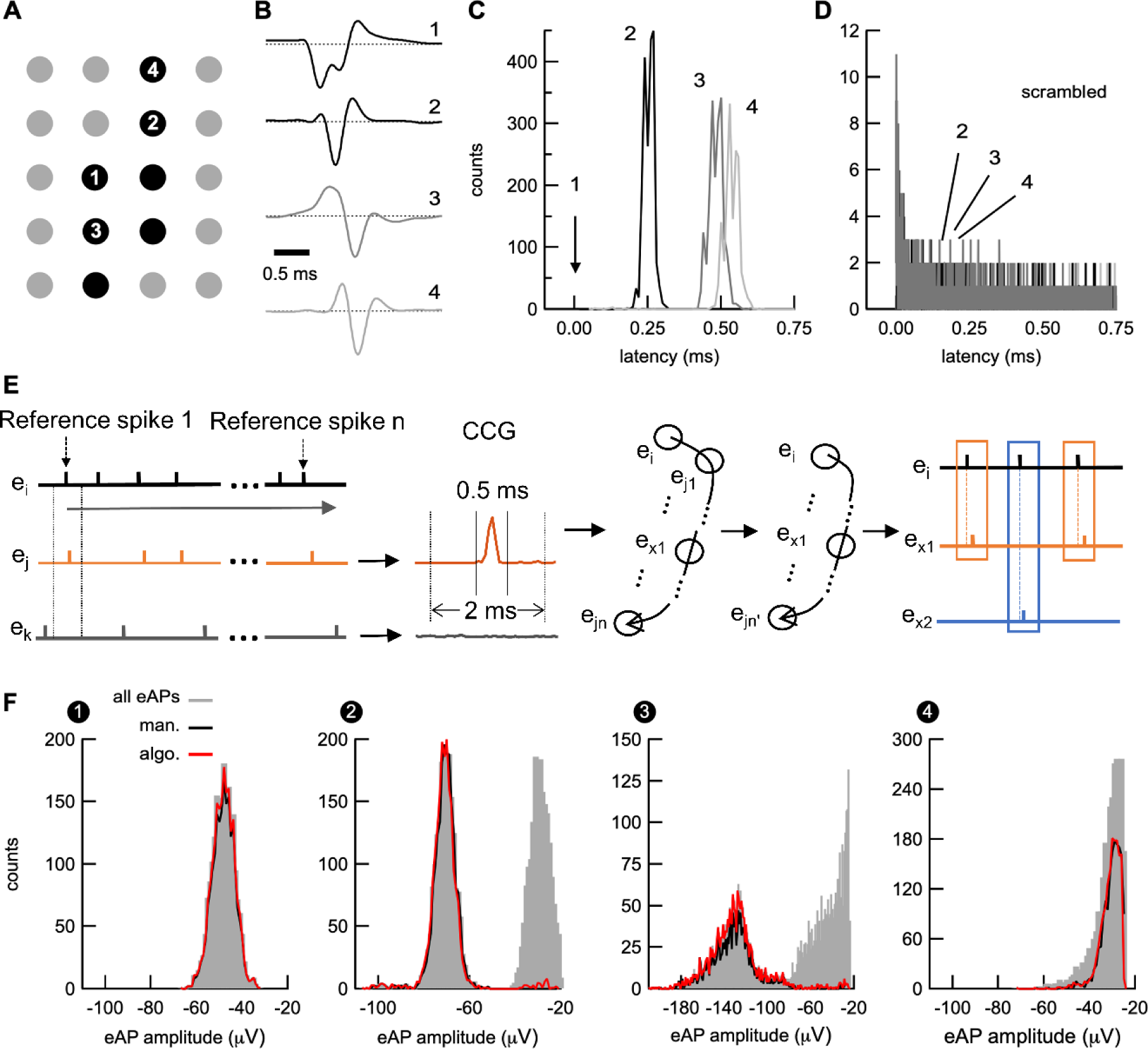
Automated detection of eAP propagation. ***A***, Action potential propagation was manually detected among electrodes indicated by black dots. ***B***, shows averaged eAP waveforms from the electrodes indicated in ***A***. The time offset of the eAP peaks, from top to bottom, illustrate propagation among the indicated electrodes. Each waveform represents the average of 90-100 sweeps and averaged waveforms were scaled to the same amplitude for the sake of clarity. ***C***, shows the eAP latency distribution between propagating eAPs at electrode 1 and electrodes 2, 3, and 4, respectively. Panel ***D*** shows the latency distribution of scrambled eAP times from ***C***, showing very few scrambled spikes occur at this short latency. Scrambled eAP latency distributions for electrodes 1-3 are superimposed. Note the difference in y-axis between ***C*** and ***D***. These features are used for automated detection, as outlined in ***E***. ***F*** compares results from manual detection of action potential propagation (black lines) and the results from our algorithm (red lines) by overlaying each on the all-points eAP amplitude histograms (grey) from the electrodes in our example. Note the small fraction of potentially mis-detected eAPs for manual and automated detection outside of the main modes in the middle histograms. This recording was done on a low electrode density array (120 electrodes; 100 μm pitch) from cultured mouse hippocampal neurons. Bin sizes in ***C*** and ***D*** were 0.1 ms. Bin sizes in ***F*** were 1 μV.

0.485 ± 0.037 ms, and 76.2% (1583/2078) of the eAPs at electrode 1 were followed by eAPs at electrode 4, with a mean latency of 0.534 ± 0.047 ms (Figure 1C). Manual assessment of eAP co-occurrences was done by comparing the eAP times in electrode 1 with spike times at other cohort electrodes and excluding inter-electrode latencies greater than 1 ms. The coefficients of variation (CV) of the latencies in this example (0.071, 0.076 and 0.088 respectively) are consistent with a high fidelity process like action potential propagation.

To test whether the latencies between eAPs at electrode 1 and eAPs at the other electrodes resulted from random alignment, we shuffled the spike times of eAPs at electrodes 2, 3 and 4 while retaining the identical inter-spike interval (ISI) distribution of those electrodes (Figure 1D). In the shuffled data, there were only 49, 69 and 92 eAPs (for electrodes 2, 3 and 4, respectively) with latencies below 1.5 ms (0.68 ± 0.44 ms, 0.76 ± 0.45 ms and 0.77 ± 0.46 ms respectively). These results suggest that the high number of eAPs at electrodes 2, 3 and 4 with sub-millisecond latency from electrode 1 were unlikely to result from random alignment of spikes and are instead consistent with action potential propagation among these electrodes.

Our automated method uses the invariant unidirectionality of eAP detection and low inter-electrode latency to isolate each electrode cohort of propagating eAPs. The sequence of eAP propagation among these electrodes uniquely identifies each unique eAP cohort. The algorithm first constructs cross-correlograms between a reference electrode and all other electrodes to find candidate constituent electrodes from each cohort of propagating eAPs (Figure 1E). In every iteration, each electrode is used as a reference electrode, to compare with all other electrodes. A collection of electrodes representing the path of eAP propagation is then generated. Within each such electrode cohort, the number of short latency (<1.5 ms) co-occurrences between eAPs at the reference electrode and eAPs at all other electrodes are compared. The maximum number of eAP co-occurrences between the reference electrode and any other cohort electrode approximates the spike times for that electrode cohort. We refer to these two electrodes as anchor points, and the electrode with the earliest eAPs in the propagation sequence as anchor point 1. A second scanning step reduces false positive electrodes by eliminating electrodes with a low fraction of co-occurrences. The algorithm outputs cohort electrodes along with statistics for each unique cohort detected, including spike times of each isolated spike train, inter-electrode latencies and number of co-occurrences between the reference electrode and other electrodes. Details of user-defined settings for detection parameters refinement, as well as criteria for selecting candidate electrodes are described in the Methods section.

The histograms in Figure 1F show the amplitudes of all spikes at electrodes 1 through 4 (grey). Superimposed on these distributions is a comparison at each electrode of the results of manually detected short latency eAPs and the isolated eAPs from automated detection (black and red lines, respectively). Automated and manual analysis revealed that the majority of eAPs in electrode 1 were followed with short latency by eAPs in the other electrodes. For automated detection, 97.1% of the eAPs at electrode 1 were followed by eAPs in electrode 2, 95.9% were followed by eAPs in electrode 3 and 78.3% were followed by eAPs in electrode 4 (red lines in each histogram). Using manual detection, these values were 93.3%, 92.0% and 76.2% respectively (black lines in each histogram). This comparison demonstrates that results from automated action potential propagation detection are at least as good by virtue of amplitude and timing information, as the far more error-prone and laborious process of manual comparison of spike times from different electrodes.

Because detection of action potential propagation confirms eAPs as resulting from a single neuronal source, the narrow eAP amplitude distribution in electrode 1, coupled with the high number of co-occurrences (>97%) detected suggests that this amplitude distribution represents eAPs from a single neuron. In contrast, the eAP amplitude distribution from electrode 2 is divided into two discrete modes. Automated detection revealed that the eAPs in electrode 2 which occurred with short latency (1980/3864) from eAPs in electrode 1 were limited almost exclusively to one mode of the electrode 2 distribution (red line). If random alignment of spikes explains the short latency between eAPs in electrodes 1 and 2, the eAP amplitudes would be evenly distributed between both modes of the electrode 2 distribution. However, these short latency eAPs were limited to an apparent single mode, consistent with a single neuronal source of those eAPs.

Propagating eAPs at electrode 3 that were isolated either by our algorithm or manually were limited to the same single amplitude distribution component (red and black lines) and overlapped each other almost uniformly. Unlike the eAP amplitude distributions from electrodes 2 or 3, the amplitude distribution at electrode 4 appears to be a single mode, part of which was excluded by the detection threshold for this electrode, which likely explains why fewer eAP co-occurrences between electrode 1 and 4 were found with either detection method. As with electrodes 2 and 3, amplitude distributions from automated and manually isolated eAPs almost completely overlap each other at electrode 4 (red and black lines). Interestingly, though the electrode 4 eAP amplitude distribution appears unimodal, our results indicate that spikes from at least 2 neurons were detected at this electrode, highlighting the unreliability of amplitude distribution shape as criteria to estimate the number of neurons at an electrode. These results demonstrate that automated detection of action potential propagation yields results that are, at minimum, as good as results from the much more time consuming and error-prone manual detection.

### Detection probability and spike train refinement

Our method for isolating eAPs from individual neurons is based on detection of action potential propagation between at least 2 electrodes. The reliability of neuronal spike trains isolated by our method is therefore sensitive to the signal-to-noise characteristics of cohort electrodes. Assume, for example, that eAPs at the first electrode have an infinitely high signal-to-noise ratio and propagating eAPs at the second electrode have a mean amplitude of 1.5 times the detection threshold, with a standard deviation that is 33% the eAP amplitude. The co-detection requirement means that in this example, the number of propagating eAPs detected will be under-sampled by at least 16% below the number of eAPs at first electrode, resulting in gaps in the spike train.

We used co-detection of eAPs in the first two anchor points to determine how eAP detection probability varies with the signal-to-noise ratio at the other cohort electrodes from our algorithm. For this analysis, the set *S* is the co-occurring eAPs between the anchor point 1 and any other cohort electrode giving the most co-occurrences. The set *S* is a proxy representing the maximum number of eAPs detected from each neuron. The detection probability for all other electrodes in the same cohort is defined as the ratio of the number of co-occurrences between eAPs at that electrode and *S* to the number of eAPs in *S*.

A map of coactive electrodes for a propagating eAP is shown in Figure 2A (black circles). Figure 2B displays the average waveforms from electrodes 1, 2 and 3.

**Figure 2.**
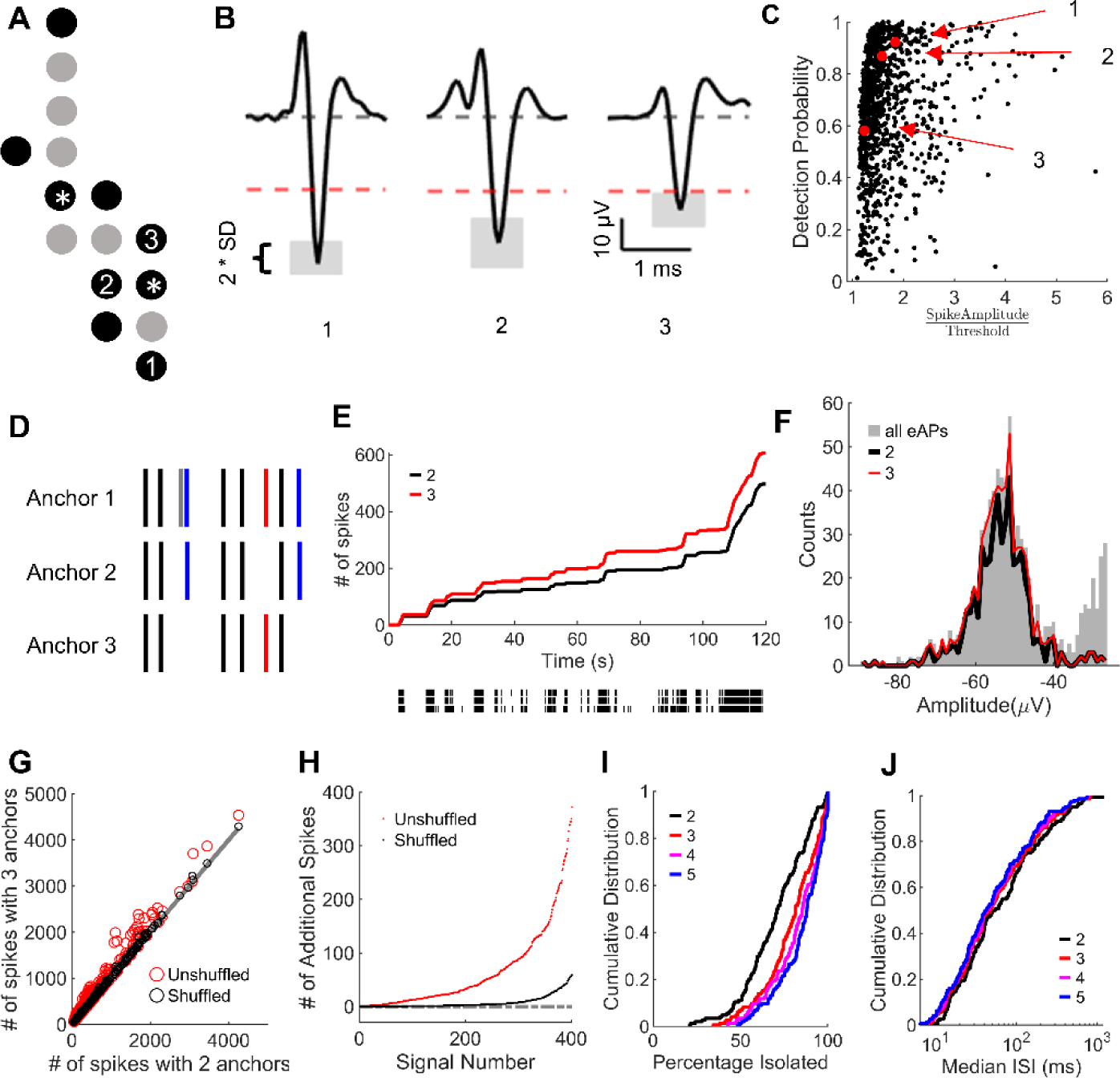
Detection probability and anchor points selection. ***A***: Map of constituent electrodes (black circles) for the subset of propagating eAP waveforms shown in (***B***). The two anchor points are marked with *. ***B***: Average of 1000 propagating eAP waveforms from electrodes 1, 2 and 3 in (***A***). Time and amplitude axes are the same for each waveform. Dashed black lines are the baseline (0 µV). Red dashed lines represent the specific spike detection threshold at each electrode. The grey areas indicate ± one standard deviation of the eAP amplitude from 1000 propagating eAPs from each electrode. ***C***, the detection probability rises steeply with increasing value of the ratio of eAP amplitude to the eAP detection threshold. This was done by using propagating eAPs in anchor points 1 and 2 as a reference. Each black dot (n = 1030) represents one constituent electrode of an eAP propagation and the red dots are results from electrode 1, 2 and 3 from ***B***. We used 750 examples of eAP propagations from 63 recordings for data in ***C***). 7 dots with a spike amplitude to threshold ratio over 6 were omitted to highlight the majority of the data. ***D***: A raster plot of the first three anchor point electrodes from one eAP propagation. In this example, black lines indicate co-occurring spikes detected by all three electrodes, blue lines show co-occurrences detected only by the first two anchors and red lines represent co-occurrences detected with the use of a third electrode. Grey line is an eAP at anchor electrode 1 with no co-occurrences with electrodes 2 or 3. ***E***, shows the comparison of the number of propagating eAPs detected during the voltage record by using 2 versus 3 anchor point electrodes. The raster plot for the three anchor point electrodes in this example is below, with anchor 1, 2 and 3 from top to bottom. ***F***, shows the histogram of all eAP amplitudes (grey) in anchor electrode 1 from ***E***. The resulting propagating eAP amplitude distributions isolated using 2 (black line) or 3 (red line) anchor points are superimposed. Electrodes from ***E*** were used in this example. ***G***, eAP propagations from 414 neurons in 63 recordings were used to compare the number of propagating eAPs isolated when using three anchor electrodes versus the number of propagating eAPs isolated using two anchor electrodes (red circles). The results of shuffling the eAP times on anchor 3 versus the number of spikes isolated with two anchors are superimposed (black circles). The slope of the grey line is equal to 1, indicating no difference between the two conditions. ***H***, compares the additional propagating eAPs detected by adding a third anchor point in shuffled versus unshuffled data (n = 414 eAP propagations). All dots are in ascending order by the number of eAPs added by including a third anchor point electrode (n = 414 eAP propagations). ***I***, Percentage of propagating spikes isolated over the total number of spikes on anchor point 1 for each eAP propagations using 2, 3, 4 or 5 anchor points. ***J***: The median ISI of the same 144 eAP propagation using 2, 3, 4 or 5 anchor points as in ***I***. The analysis in ***I*** and ***J*** is done with 144 eAP propagations with at least 5 cohort electrodes, in 63 recordings. All data in this figure were from recordings done on low electrode density arrays (120 electrodes; 100 μm pitch) from cultured mouse hippocampal neurons.

Waveforms are ordered from left to right by their eAP amplitude/detection threshold ratios. As expected (Poor, 1994), the detection probability rises steeply with increasing value of the eAP amplitude/detection threshold ratios (Figure 2C). This plot includes data from 1030 electrodes of 750 unique propagating eAPs in 63 recordings from neurons on low density arrays (120 electrodes; 100 μm pitch). Red dots are values from the example eAP waveforms (Figure 2B). Failures of propagation cannot be ruled out as occasionally contributing to failures to detect eAPs at cohort electrodes (Grossman et al., 1979; Soleng et al., 2004). However, given the high coefficient of variation of the amplitude/threshold ratio (1.73± 0.91, n = 1030) and the high fidelity of action potential propagation (Cox et al., 2000; Raastad and Shepherd, 2003), the occasional absence of expected propagating eAPs at a cohort electrode (Figure 2C) is consistent with a failure of spike detection. The results reported here with automated eAP propagation detection recapitulate results previously obtained manually (Tovar et al., 2018).

These results demonstrate that to maximally represent spike trains from isolated neurons, selecting coactive electrodes with high detection probability is crucial. The relationship between eAP detection probability and the eAP amplitude/threshold ratio may result in under-sampling spikes, thus creating gaps in the spike train. However, if detection failures at any electrode occur randomly, then detection failures at cohort electrodes would not be expected to occur simultaneously (Figure 2D). To minimize gaps in the spike train resulting from failures of spike detection, we examined whether adding anchor points improves the results of the automated detection of eAP propagation algorithm. The algorithm first identified constituent electrodes of eAP propagation. Co-occurrences between eAPs at the first 2 anchor points *(S*) are used to approximate the number and timing of propagating spikes. We used the same 1.5 ms detection limit following the eAP at anchor point 1. Cohort electrodes were ordered based on the number of co-occurrences with anchor point 1. The electrode with the most co-occurrence is designated anchor point 2. The propagating eAP spike times of anchor point 1 are the best representation of the spike train from that neuron, due to the high signal-to-noise ratio near action potential initiation (Bakkum et al., 2013). We tested the effect of adding additional anchor points, from most to least co-occurrences with anchor 1. Co-occurrences between additional anchor points and anchor point 1 are added to *S* to comprise an amended set of propagating eAPs.

The schematic (Figure 2D) illustrates the effect of using two versus three anchor points. In this example, if propagation is defined by eAP co-occurrences at anchor 1 and anchor 2, then 7 eAPs will be isolated. Including anchor 3 increases the total to 8 eAPs. Figure 2E compares the effect of using 2 versus 3 electrodes to reduce artifactual gaps in the spike train. Using 3 anchor points results in more isolated eAPs (Figure 2E) compared to using 2 anchor points, as shown by the progressively greater increase in number of propagating eAPs during the recording. The amplitude distribution of the extracted eAPs from anchor 1 from this example (Figure 2F) shows that spikes isolated with the addition of a third electrode occur in the same amplitude mode, indicating that those eAPs are from the same single neuronal source. At the anchor point 2 electrode, the mean amplitude of propagating eAP was -89.7 ± 10.0 μV and the threshold was -24.1 μV which should allow us to isolate almost all propagating eAPs (Figure 2C). However, including propagating eAPs from anchor point 3, (-35.0 ± 4.9 μV mean amplitude; -24.8 μV threshold) still increased the total number of propagating spikes that were isolated by our algorithm, demonstrating how gaps in the spike train were filled by increasing the number of anchor points when possible.

Using more than 2 anchor points reduced the number of undetected propagating eAPs but could also introduce artifactually detected eAPs to the spike train. To test the extent to which such co-occurrences added by a third anchor point represent random eAP alignment between anchor point electrodes, we shuffled the timing data at the third electrode, then determined the number of eAPs added to the propagation spike train (Figure 2G). Data shuffling was done by retaining the ISI distribution but shuffling the spike time. This was done with 414 eAP propagations that had at least three constituent electrodes, from the same 63 recordings. If inclusion of a third anchor point produces no increase in the spike total, then all points will lie on the line with the slope equal to 1 (grey line). The number of spikes added when including a third anchor point increased more in the unshuffled spike set (red dots) than in the shuffled spike set (grey dots). A direct comparison of the number of additional spikes with or without shuffling (Figure 2H) indicates that shuffling only occasionally increased the number of spikes, whereas using a third anchor point almost always added eAPs to each spike train (Figure 2H). For example, in 138/414 neurons, using 3 anchor points increased the number of propagating eAPs by at least 20% compared to using 2 anchor points. In contrast, shuffling the spike times at the third electrode resulted in a 20% or greater increase in additional eAPs in only 4/414 neurons. In unshuffled data, using 3 anchor points resulted in a 30% or greater addition of propagating eAPs in 86/414 neurons whereas there was never any increase in the number of spikes greater than 30% in the shuffled data. These results demonstrate that random alignment of eAPs is not a significant source of spikes added when using a third anchor point.

Increasing the number of anchor points produce significant refinements in spike rate detection of the propagating eAPs in our data. We therefore examined the result of adding additional electrodes in the same data set by comparing the results of using 2, 3, 4 or 5 anchor points. Increasing the number of anchor points results in a gradual increase in the percentage of propagating eAPs isolated (Figure 2I). In this data set (144 eAP propagations with at least 5 cohort electrodes) the increase from 2 and 3 anchor points diminishes when adding 4 or 5 anchor points. The median ISIs from the same 144 eAP propagations comparing the number of anchor points results in relatively minor differences above 3 anchor points (Figure 2J). For different use cases, the number of anchor points in our algorithm is a user-defined variable. Our results demonstrate that most propagating eAPs can be found with just two anchor points. However, using more anchor points refines the extracted spike train, resulting in fewer gaps due to detection failures, with diminishing differences above 3 anchor points in this data set.

### Automated detection of propagating eAPs is insensitive to frequency-dependent changes in eAP waveform

Transmembrane action potentials in many types of neurons show frequency-dependent changes in amplitude or spike width (Jackson et al., 1991; Liu et al., 2017). Frequency-dependent changes are reflected in the extracellularly recorded action potentials from these neurons (Buzsáki et al., 1996; Fee et al., 1996). Spike sorting routines could interpret frequency-dependent eAP waveform heterogeneity from single neurons as action potentials from multiple neuronal sources. The problem for spike sorting resulting from heterogeneity in action potential waveform in single neurons has been discussed elsewhere (Quirk et al. 1999; Stratton et al., 2012). Unlike spike sorting routines, detection of action potential propagation reliably isolates spikes from single neurons when frequency-dependent changes in eAP waveform occur (Figure 3). This example illustrates that a burst of propagating action potentials in one constituent electrode (Figure 3A) results in a frequency-dependent decrease in spike amplitude (Figure 3B) and an increase in spike width (Figure 3C). Note the change in the peak-to-peak latency of eAPs that occurred following ISIs ≥ 500 ms, compared to the latency for eAPs following ISIs of ≤ 30 ms (brackets, Figure 3C). This difference likely reflects decreased Na^+^ channel availability at high spike frequency (Colbert et al., 1997). As shown, our method of detecting action potential propagation isolates eAPs from single neurons at inter-spike intervals as short as 2 ms (Figure 3D), just above the refractory period frequency cutoff we used for spike detection. The cumulative distribution of inter-spike intervals (ISI) shows that in this recording, 64% of all the spikes (open grey circles) occurred following ISIs from 30 to 500 ms. The plot of eAP amplitude as a function of ISI shows the amplitude variability in this intermediate frequency domain (Figure 3E).

**Figure 3.**
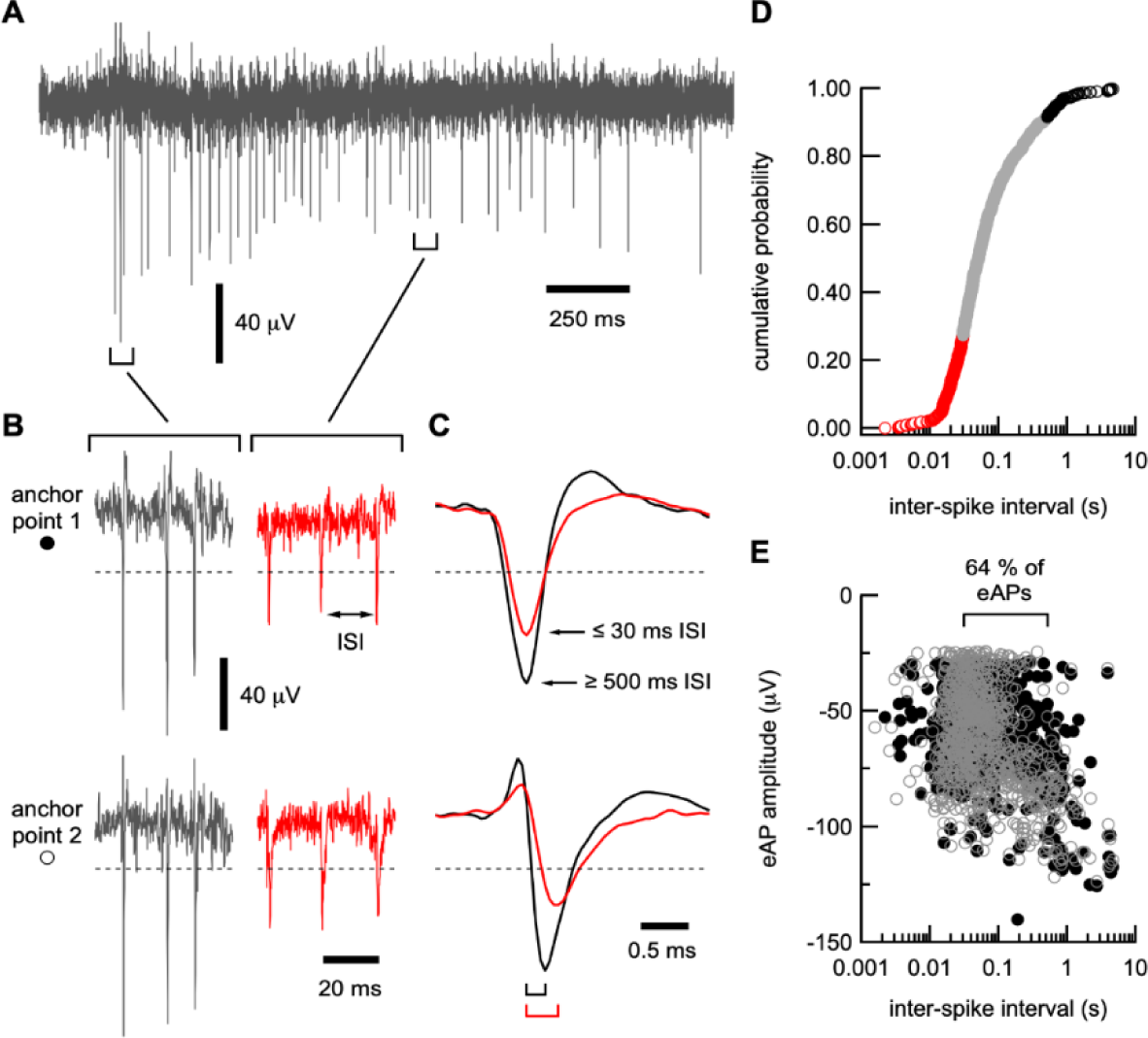
Frequency-dependent changes in eAP waveform in single neurons revealed by automated eAP propagation detection. ***A*,** voltage record from anchor point 1 electrode showing a frequency-dependent reduction in eAP amplitude. For clarity, only one of two anchor point electrode voltage traces from this propagating eAP is shown here. This burst was preceded by an ISI of >2 s. ***B*,** eAPs in the anchor point 1 and anchor point 2 electrodes, shown at the beginning of the burst (grey traces) and in the middle of the burst (red traces). ***C*,** for the propagating eAP from this neuron, we superimposed the average of 50 eAPs that were preceded by ISIs of ≥ 500 ms duration (black traces) on 50 averaged eAPs that were preceded ISIs of ≤ 30 ms. Note the frequency-dependent difference in amplitude and spike width of eAPs from each anchor point electrode. Amplitude scale is the same in ***B*** and ***C***. Dashed lines in ***B*** and ***C*** indicate the eAP detection threshold for each electrode. The averaged waveforms for anchor point 2 eAPs are time referenced to propagating eAPs from electrode 1. Brackets below the waveforms in ***C*** indicate the inter-electrode latency for long (black) and short (red) ISIs. ***D*,** the cumulative distribution of ISIs from this neuron show large distribution in time. The eAPs that occurred ≤ 30 ms are shown by open red circles, eAPs that occurred ≥ 500 ms are shown by open black circles. Outside of these spike frequency ranges lies the majority of ISIs (open grey circles). ***E***, the relationship between eAP amplitude and ISI in this neuron, showing that the 64% of eAPs occurred after ISIs between 30 and 500 ms for anchor point 1 (closed circles) and anchor point 2 (open circles). This highlights how eAPs kinetics and spike train data from single neurons can be extracted in spite of the frequency-dependent changes in spike morphology.

For example, for the propagating spikes in the anchor point 1 electrode (closed circles) that follows intervals between 30 to 500 ms had a mean amplitude of -54.9 ± 15.8 μV (n = 552). At the anchor point 2 electrode (open circles) in the propagation pathway from this example, the mean amplitude of propagating spikes (from anchor point 1) occurring within intervals of 30 to 500 ms was -61.7426 ± 21.5336 μV (n = 554), mirroring the amplitude variability in eAPs in the anchor point 1 electrode. Because eAP morphology is expected to vary with inter-spike interval (Buzsáki et al., 1996), the variability of spike shapes in this large frequency range may present as eAPs from multiple units for spike sorting. However, our algorithm detected action potential propagation in this neuron across more than three orders of magnitude of inter-spike intervals (Figure 3D, E).

### Automated detection of propagating eAPs is insensitive to experimentally-induced changes in waveform shape

Many experimental manipulations change the eAP waveform, either by modulating intrinsic behavior of ion channels underlying transmembrane conductances or by directly targeting the ion channels active during action potentials. Spike sorting routines which cluster eAPs on the basis of shape are challenged by common experimental manipulations that produce changes in the waveform shape, leading to failures in correctly identifying eAPs from the same neuronal source across changing experimental conditions (Hilgen, 2019). However, because our method of isolating spikes only depends on the sequence of propagating eAPs at cohort electrodes it is insensitive to changes in eAP waveform.

Our algorithm isolates eAPs from single neurons between experimental conditions that change eAP waveform shape, as shown by comparing the effects of the K^+^ channel antagonist 4-AP (100 μM) on propagating eAP waveforms in different anatomical places from the same neuron (Figure 4A). Overlaying control eAPs (black traces) and eAPs in 4-AP (red traces) highlights changes due to K^+^ channel antagonism in this cultured neuron. Note the heterogeneous effects of 4-AP on eAP waveforms at different points in the propagation pathway from this neuron, including increased duration (panels 1 and 2), decreased eAP amplitude (panel 3) and reducing the repolarization phase (panels 1 - 4). These results illustrate the difficulty in predicting how any drug will affect eAP waveform. Waveform heterogeneity is likely due to how capacitive and resistive components of the eAP waveform combine and are affected by the extracellular electrical field (Buzsáki et al., 2012). Figure 4B shows the multimodal amplitude distribution in electrode 4 of the propagating action potential from Figure 4A, in control (left) and during 4-AP application (right). The amplitude distribution of the propagating eAPs at electrode 4 are indicated in red. In control condition, eAPs isolated from this neuron constitute 37.3% of the eAPs at this electrode (778/2083) which increased to 49.1% (1278/2604) of the eAPs in 4-AP (in a 3-minute recording). Details of how experimental manipulations affect the spike trains from these neurons are seen in the ISI distributions from these data (Figure 4C). Solid lines indicate the ISI for all spikes at electrode 4 and dashed lines indicate the ISI of the propagating eAPs, showing in this example no systematic change in the spike train resulting from 4-AP application (Figure 4C).

**Figure 4.**
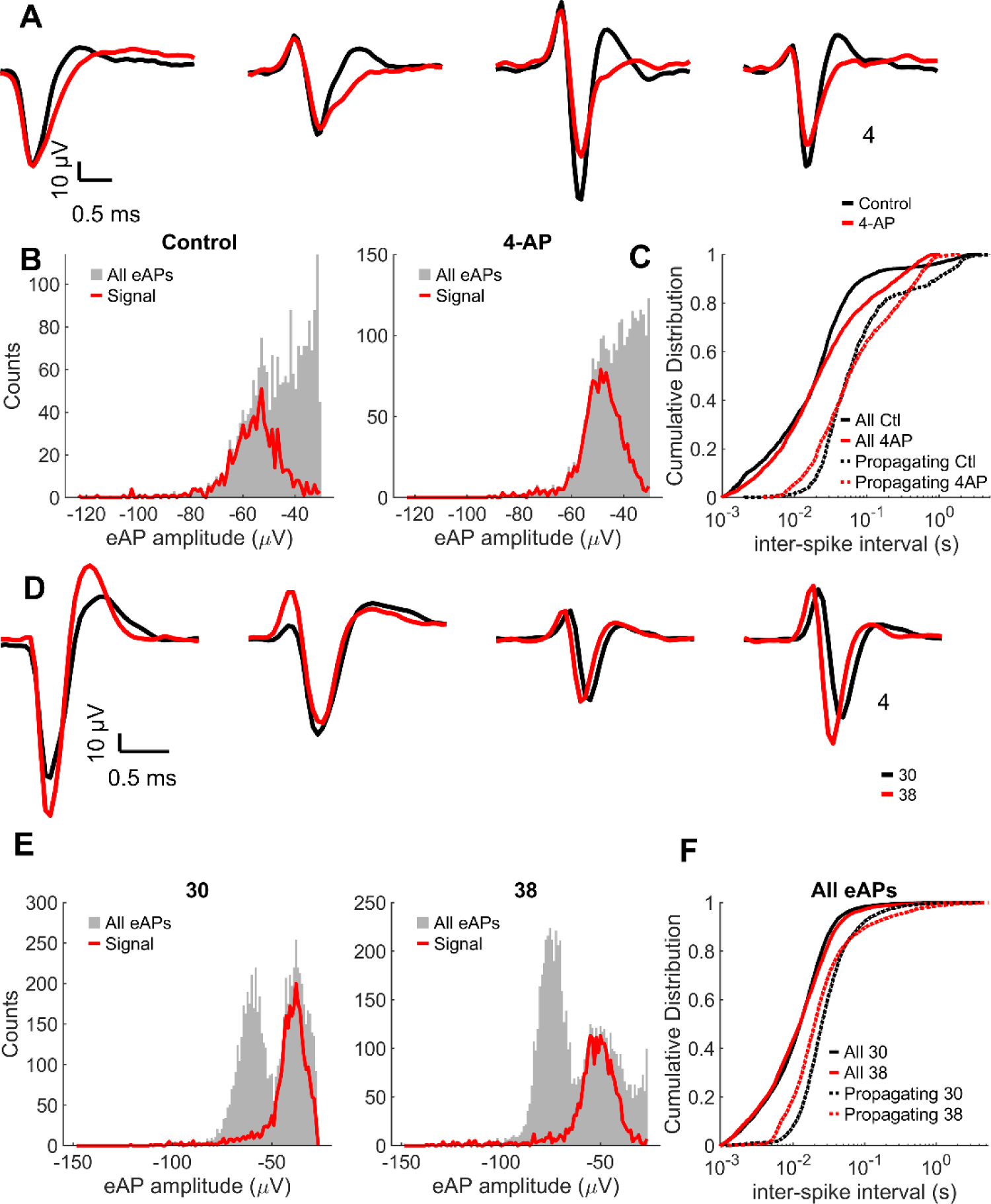
Automated detection of eAP propagation during experiments that change eAP waveform. ***A***, the averaged propagating eAP waveforms from four constituent electrodes in control condition (black traces) and in 100 µM 4-AP (red traces) are superimposed, illustrating the effect of 4-AP on eAP waveforms at different points in the propagation pathway. ***B***, single neuron propagating eAP amplitude distribution in control and 4-AP, compared with the all-points amplitude distribution in electrode 4. ***C***, the ISI distribution of all eAPs in electrode 4 versus the ISI distribution of only the propagating eAPs, comparing control and 4-AP. ***D***, in another experiment, the averaged propagating eAP waveforms from 4 constituent electrodes are superimposed to compare waveforms shape in recording at 30°C (black traces) versus recording at 38°C (red traces). Averaged waveforms for electrodes 2 - 4 are time referenced to electrode 1. Note the leftward shift of the eAP peak in the red traces relative to black, consistent with the temperature induced increase in propagation speed. ***A*** and ***D*** illustrate that experimentally induced changes in eAP waveform do not affect the ability of our algorithm to detect spikes from the same neuron. ***E***, single neuron propagating eAP amplitude distribution at 30°C versus 38°C, each superimposed on the all-points amplitude distribution in electrode 4. Note the increase in eAP amplitude at higher temperature. ***F***, the ISI distribution of all eAPs in electrode 4 versus the ISI distribution of only the propagating eAPs, comparing 30°C and 38°C. Data in this figure were taken from recordings done on low electrode density arrays (120 electrodes; 100 µm pitch) using cultured mouse hippocampal neurons.

In another experiment, varying the recording temperature from 30°C (black traces) to 38°C (red traces) also changed the morphology of propagating eAP waveforms at four constituent electrodes from a single neuron (Figure 4D). Higher temperature altered the spike width and amplitude (panel 1, 4) and the capacitive component of the eAP (panel 2). Note the leftward shifts of eAPs at higher temperature (panel 3-4), reflecting faster propagation at higher temperature (Chapman, 1967; Franz and Iggo, 1968). Regardless of changes in waveform morphology, our algorithm successfully identified propagating eAPs at the same four constituent electrodes. The amplitude histograms of propagating eAPs at electrode 4 from the temperature experiments (Figure 4E) are superimposed on the amplitude histogram of all eAPs at that electrode. For this neuron, propagating eAPs were 42.1% (2849/6768) of all eAPs at electrode 4 at 30°C. Propagating eAPs decreased to 32.9% (2234/6787) of all eAPs at this electrode at 38°C. Propagating eAPs isolated by our algorithm were limited to an apparent single mode of the multimodal, all-point amplitude distributions from these electrodes, even when experimental conditions changed the eAP amplitude distribution (Figure 4B, E).

Unpredictable changes to the eAP amplitude distributions such as these (Figure 4B, E) may result in unreliability of the results of spike sorting. The leftward shift in the ISI distribution from isolated eAPs in this data (red versus black dashed lines) clearly illustrates the effect of higher temperature increases eAP propagation speed (Figure 4F). The temperature effect is obscured by overlapping ISI distributions of all spikes in electrode 4 (Figure 4F). These results demonstrate that isolating spikes from single neurons based on action potential propagation is insensitive to changes in the eAP waveform and that our algorithm reliably isolates spiking in single identified neurons during changing experimental conditions.

### Our algorithm works with multiple extracellular recording platforms

Our algorithm works with any type of multi-electrode spike trains, with no upper limit on the number of electrodes. The time complexity of our algorithm is proportional to:

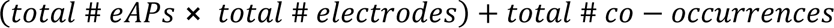

Multiple propagating eAPs are readily detected with higher density planar CMOS arrays (26,400 electrodes; 17.5 micron pitch). An example of a subset of eAP propagations detected from a 21 x 24 electrode section of CMOS array containing 330 active electrodes, and their footprints generated by signal averaging based on anchor point 1 for each eAP propagation, are shown (Figure 5A). For ease of visualization, not all examples of propagating eAPs are displayed. Each waveform represents an electrode from which super-threshold eAP components were recorded (Figure 5A). As we’ve done here, the results of our algorithm could be used for refinement of axonal propagation pathway for each detected signal (Bakkum et al., 2013).

**Figure 5.**
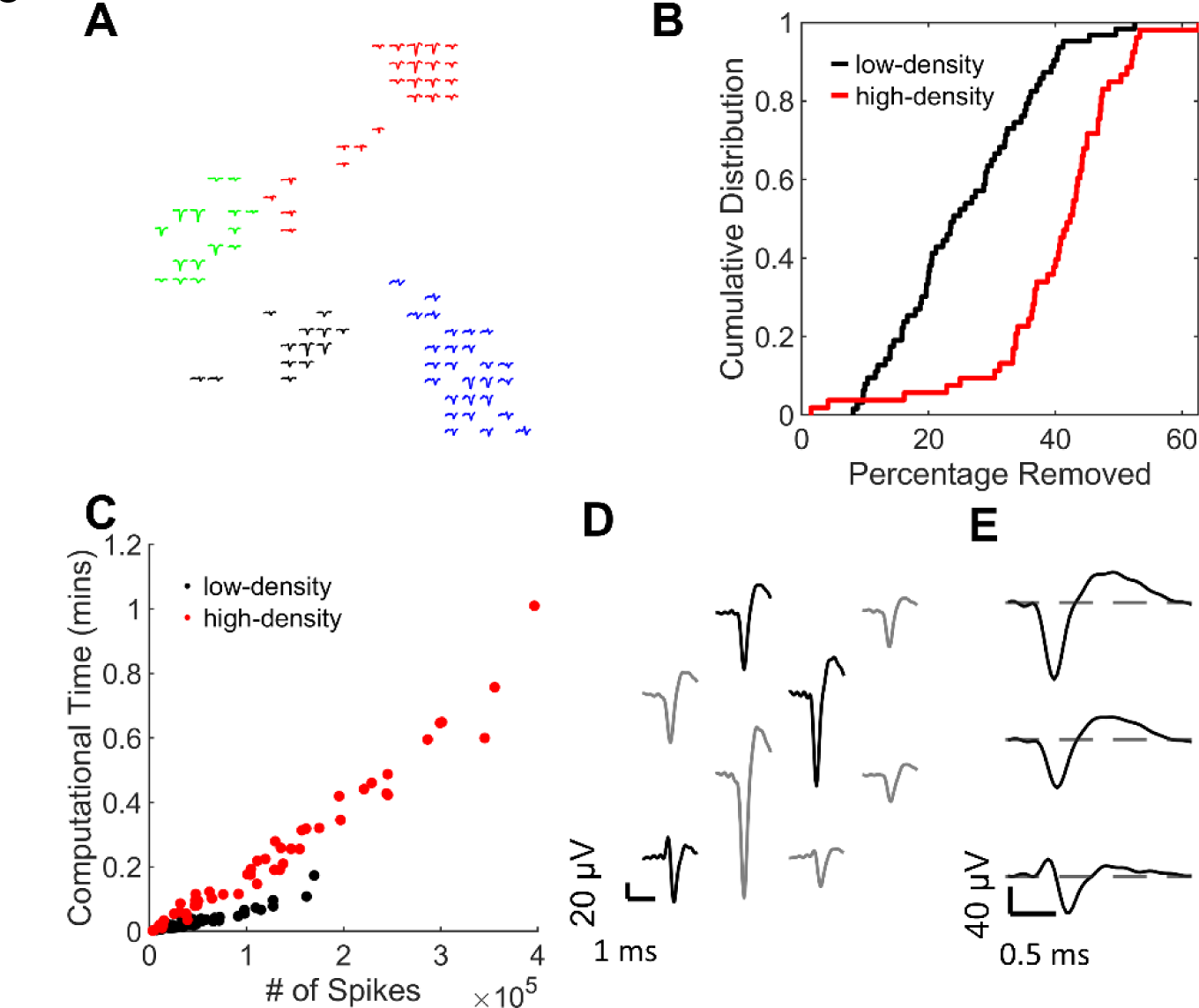
Automated detection of eAP propagation handles data from a range of recording platforms. ***A***: a subset of eAP propagations detected on the CMOS array (26,400 electrodes, 17.5 μm pitch). Each color is the footprint of propagating eAPs in one neuron on this 21 electrode by 24 electrode section, with 330 active electrodes. Each footprint is generated by signal averaging based on anchor point 1 for propagating eAPs in each neuron. ***B***. The percentage of ’redundant’ spikes eliminated in each recording from low density arrays (black) or CMOS arrays (red). ***C***: Computational time of the algorithm versus the number of spikes with the data from 63 low-density recordings (black) and 53 high-density recordings (red). ***D***: A footprint of an eAP propagation detected on a Neuropixels recording. The footprint is generated by signal averaging based on anchor point 1 for each propagating eAPs. Black lines are the same waveforms shown in **(*E*)**. Note the time offset from top to bottom in **(*E*)**, indicating propagation among the three electrodes in three-dimensional tissue from this organoid. Low density arrays have 120 electrodes with a 100 μm pitch. High density CMOS arrays have 26,400 electrodes with a 17.5 μm pitch. Recordings on low and high density arrays were done using cultured mouse hippocampal neurons. The Neuropixels probe was used to record eAPs from a cerebral organoid derived from human iPS cells.

Propagating eAPs from single neurons are often detected at multiple electrodes, especially in high-density arrays. The spikes at these electrodes are redundant because a neuron’s spiking pattern can be represented by eAPs at anchor point 1. The number of constituent electrodes artifactually multiplies the number of spikes from a neuron. In the case of spike sorting, redundant eAPs such as these could be mistakenly isolated as independent units unless they are removed from the data record. Higher electrode density results in greater number of redundant eAPs; the mean number of cohort electrodes for each eAP propagation on high-density CMOS arrays was 14.1 ± 8.9 electrodes (n = 1606 neurons). In contrast, each eAP propagation recorded on low-density arrays was detected by 3.5 ± 2.0 electrodes (n = 750 neurons), demonstrating that the extent of the spike redundancy of propagating eAPs varies with recording platform and electrode density. To demonstrate this explicitly, we removed propagating eAPs from all cohort electrodes except for anchor point 1 in multiple recordings from both platform types and plotted the distribution of the percentage of spikes removed from both platforms (Figure 5B). In 6/63 recordings using low-density arrays and 33/53 recordings using the high-density arrays, 40% or more of the eAPs in the entire data record could be removed due to redundancy. Similarly, 40/63 of the low-density recordings and 50/53 of the high-density recordings have at least 20% redundant spikes. The relatively high number of cohort electrodes for each eAP propagation in high-density arrays could improve spike train refinement due to more choices for selecting anchor point electrodes with high-signal-to-noise characteristics. These data are from 2-minute-long recordings from low- and high-density arrays. For ease of data acquisition, high-density arrays (26,400 electrodes) were segmented into 32 blocks. Each block was recorded independently because only 1024 electrodes can be recorded simultaneously. This produced 53 recordings from high-density arrays with at least 1 example of eAP propagations, which we compared to 63 recordings in low-density arrays. In arrays with addressable electrodes, our algorithm maximizes the recording efficiency by identifying anchor points for individual neurons.

Simultaneous recording of multiple ground-truth validated neurons can then be done in subsequent experiments using the minimum number of electrodes needed to isolate propagation eAPs for each neuron.

We compared the computational time required for our algorithm to output results from the low- or high-density arrays described above (Figure 5C). The specification of the server on which this analysis was done is described in Methods. For a recording with 100000 spikes, the approximate computational time was 3.5 seconds for the low-density arrays we used, and 11 seconds for our high-density arrays. In general, with the same number of spikes, the computational time on a low-density array is less than on a high-density array. This is due to the increase in pairwise electrode comparisons with increasing electrode numbers.

We have shown that eAP propagation can be detected from neurons grown on planar arrays. The three dimensional orientation of axonal process in intact tissue theoretically makes detection of action potential propagation less likely. We examined whether our algorithm could be used to detect propagation in recordings done on a whole cerebral organoid, using a Neuropixels probe. We identified 15 unique neurons based on automated detection of eAP propagation from a Neuropixels recording of 384 electrodes. An example of an eAP propagation detected is shown in Figure 5D. The footprint of the eAP propagation was generated by signal averaging based on anchor point 1. Figure 5E shows the same waveforms from the ones in black in Figure 5D. Note the time offset from top to bottom from these electrodes. These data demonstrate that with high density arrays, isolating eAPs from single identified neurons in 3 dimensional neural tissue is possible with automated detection of action potential propagation.

## Discussion

### Spike trains from single identified neurons with multi-electrode arrays

Extracellular recording has been used for decades to non-invasively study excitable cells in a range of experimental contexts (Hubel and Wiesel, 1963; Katz and Miledi, 1964; Wainger et al., 2014; Fernandez-Ruiz et al., 2019). Using extracellular recording to understand how neural networks integrate input, convert perception to electrical signal or respond to pharmacological or experimental manipulations requires unambiguous knowledge of the cellular source of each action potential in the data record. Discriminating the cellular source of any eAP becomes complicated because action potentials from multiple neurons can be detected by single electrodes (Figure 1F, 4B, 4E). This is especially true when neuron density is high. Post-acquisition data processing from contemporary MEAs requires automated approaches to identifying eAPs from single neurons.

Spike sorting routines are based on assumptions regarding how characteristics of eAP waveform heterogeneity correspond to the source neurons where those eAPs originate. Ground truth validation of spike sorting is not routinely available but has been achieved in technically challenging experiments pairing extracellular MEA recording with intra- or juxtacellular recording (Yger et al., 2018; Jackel et al., 2017; Anastassiou et al., 2015). As previously shown (Bakkum et al., 2013; Tovar et al., 2018) action potential propagation is reflected by two or more MEA electrodes with eAPs that repeatedly occur with short latency, in the same sequence (Figure 1B, C). Characteristics of propagating eAPs identify them as being from single neurons, in an analogous way that paired recording has been used to unambiguously isolate spikes from individual neurons. When electrodes detect eAPs from multiple neurons, isolation of propagating eAPs by our algorithm labels spikes from each propagation as resulting from single neurons, even in a background of spikes from other cells. Multi-point detection inherent in our algorithm reinforces the reliability of our results. Isolation of eAP by using propagation makes no assumptions about waveform shapes. Our method of unambiguously isolating eAPs from multiple different neurons in each recording increase the utility of multi-electrode arrays for high content screening applications and could eventually be adapted for use *in vivo*, for example in retina (Li et al., 2015).

### Limitations of automated detection of action potential propagation

Because the method we outline here uses action potential propagation to isolate eAPs from individual neurons, a minimum of two electrodes are required for propagation detection. Propagation from some neurons may be undetectable due to their axons being outside the detection radius of any electrode. This is especially true for lower electrode density arrays. However, denser arrays of electrodes will result in increased detection of propagation. In the case of high electrode density arrays implanted in the human brain organoid (Figure 5D, 5E), the blind insertion of the shank array and the unknown cellular anatomy likely contributed to the limited detection of propagating action potentials. With respect to the in vivo application of our algorithm, detection of propagating eAPs may be facilitated in experiments where shank electrode arrays are inserted parallel to fiber tracts in white matter, for example.

The input of our algorithm is spike trains that have been detected by other means.

Sampling the true spike train from any neuron is always limited to signal-to-noise considerations. Our algorithm depends on co-detection of eAPs at two cohort electrodes and therefore increases the risk of under-sampling the true spike train when the signal-to-noise ratio at any anchor point is low. Because our algorithm always uses the cohort electrode with the earliest eAPs in the propagation sequence as anchor point 1 and indexes the propagating eAPs at all other cohort electrodes to the earliest eAPs, detection accuracy is limited by the signal-to-noise at this electrode. The earliest electrode often includes eAPs with waveforms consistent with them arising from at or near the axon initial segment (AIS; 1B, 4A, 4D). Because the transmembrane conductance at the AIS is large (Kole et al., 2008; Bakkum et al., 2013), the signal-to-noise of these eAPs is typically quite high. However, propagation detection is compromised the lower the signal-to-noise ratio is of other cohort electrodes (Figure 2C).

We circumvented some signal-to-noise limitations of spike detection by designing into our algorithm a user-defined option to include spike trains from cohort electrodes other than anchor points 1 and 2 (Figure 2). Detection failures at the anchor point 2 electrode could be compensated for by increasing the number of electrodes for propagation detection. Doing so revealed more propagating spikes (Figure 2E, F).

However, using propagating eAPs from other cohort electrodes refines the spike train (Figure 2E) but the increased comparisons between spikes increases the risk of false positives due to random alignment (Figure 2H).

Repeated co-occurrence of eAPs is the detection criteria for isolating propagating action potentials from different source neurons. Artifactual alignment of spikes that are not reflective of propagation at other electrodes can be mistaken as propagating eAPs.

This concern increases when the spike rate at any electrode is high. However, as we show (Figures 1D, 2G, 2H), the number of artifactually isolated propagating eAPs due to randomness was minimal under our recording conditions. Propagating eAP detection can be affected by additional thresholding based on the standard deviation of inter-electrode latencies which gives rise to the CCG. Propagating action potentials have low variability of the inter-electrode latency. Thus, eAPs at other electrodes with latencies that are at the edges of the latency distribution are more likely to represent noise. Supplementary Figure 1 displays an example of additional thresholding based on the standard deviation of inter-electrode latency.

Signal quality affects propagation detection in other ways. For example, our algorithm uses parameters such as the shape of the CCG and the number of propagating eAP co-occurrences (see Methods) to identify cohort electrodes. When detection failure varies between recordings, such as during experimentally induced changes in propagating eAP waveforms (Figures 3, 4), these parameters can be affected and can result in identification of propagating eAPs with different numbers of cohort electrodes. However, the majority of spike train data from propagating eAPs in each neuron is usually represented by 2 to 3 electrodes (Figure 2I, J), within the limits of detection. Analytically, this outcome matters, for example, when trying to increase the anatomical detail of the propagation pathway. Changing the spike detection threshold could reveal such cases.

### Future Directions - complementary use with traditional spike sorting routines

Use of contemporary MEAs requires automated approaches for post-acquisition data handling, due to the large number of sensors that each acquire data at the high enough acquisition rates to represent eAPs. To understand how any experimental manipulation affects spike rate, extracellular recording on the scale possible with contemporary MEAs requires automated methods of isolating spikes into different sources. However, sorting routines that are untested against contextually relevant ground truth could produce erroneous results in the absence of other information. In the absence of ground truth, even the most sophisticated sorting routines are unable to validate how to correlate spikes at single electrodes in experiments that significantly change the eAP waveform (Hilgen, 2019). Our approach to isolating eAPs from different source neurons based on detection of eAP propagation, routinely provides contextually relevant ground truth and thus could be used to test spike sorting routines in every experiment in which propagation is detected. For example, at each electrode, eAPs assigned to individual units by spike sorting routines could be tested against eAPs at that same electrode that were isolated as part of a propagation cohort. The extent of overlap between results from a sorting routine and the results from our method assesses the accuracy of a sorting routine (Tovar et al., 2018).

Isolation of eAPs by automated detection of action potential propagation performs the same task as traditional spike sorting. The characteristics of action potential propagation themselves validate that eAP cohorts isolated by this method are from individual neuronal sources. Thus all spikes isolated by our method can be ignored by traditional spike sorting routines. However, the characteristics of propagation detection mean that the fraction of spikes in any data set that can be isolated by this method will depend on factors such as electrode and neuron density. For example, densely packed electrodes more readily detect propagation than more widely spaced electrodes (Figure 5B). In intact tissue, positioning electrode arrays within fiber tracks or other neuronal propagation pathways should facilitate propagation detection. Data acquisition in the range of experimental contexts in which MEAs are used means that automated detection of action potential propagation and traditional spike sorting routines are complementary. A workflow in which spike train data is processed by our algorithm, followed by a traditional spike sorting method, may reduce the vulnerability of the remaining ensemble spike train to be miss-sorted. This also increases the reliability of eAP isolation of the entire data record because of the empirical isolation of eAPs with our algorithm as the first step.

Sorting eAPs into source neurons based on action potential propagation avoids limitations of waveform-based sorting. Because eAPs are isolated into unique sources based on the propagation pathway rather than waveform, drug development assays based on extracellular recording on high throughput MEA platforms become unconstrained by ambiguity introduced when drug application changes spike waveform (Hilgen, 2019).

Other experimental manipulations that change eAP waveforms can also be unambiguously studied with the implementation of our algorithm because the response of each identified neuron can be tracked among various conditions. When eAPs are unambiguously assigned to a unique neuronal source, those eAP times are an index against which eAPs from other neurons can be timed, for example to identify synaptic coupling among small numbers of neurons (Guzman et al., 2021) or to assess how genetic background affects synaptic coupling, for example. Further development of our methods can start to bridge the gap between rapidly evolving MEA technology and science that use of such technology can reveal.

## Materials and methods

### Propagating eAP detection algorithm

The input of the algorithm is spike times on all electrodes in a recording. The first step in our algorithm is to select candidate electrodes that could be cohort electrodes of an eAP propagation. Let *E* denote the total set of electrodes in an array and *n* denote the total number of electrodes in *E*. Each electrode in *E* with an average spiking frequency higher than *v1* Hz is used as a reference electrode (*ei*) to compare with all other electrodes *ej* ∈ *E* (*j* = 1, 2, …, n). This threshold value for spiking frequency *v1* ensures a minimum number of spikes on each reference electrode for the analysis in the next steps. *v1* is a user-definable parameter. In this paper, we chose *v1* to be 1Hz. We used this frequency threshold because most of our recordings were 2- or 3-minutes duration. In longer recordings, *v1* can be adjusted to a lower value. We also included an option to threshold for the minimum number of spikes in total instead of the spiking frequency.

Cross-correlograms (CCGs) using a 1.5 ms window before and after reference time-points with a bin size of 0.05 ms were then constructed for all (*ei*, *ej*) pairs. A sharp peak in a CCG indicates a strong correlation of spike times between *ei* and *ej*. To quantify sharp peaks, let *n1* denote the largest sum of counts in any 0.5 ms moving window in the CCG and *n2* denote the sum of counts of a 2.05 ms window centered at the bin with the largest sum (if the largest sum is found in the first 1 ms or the last 1 ms of the CCG, take the sum of the counts of the first 2.05 ms window or the counts of the last 2.05 ms window as *n2*). The bigger the ratio *n2/n1* is, the sharper the peak is. A customizable variable *v2* is used to threshold the lower bound of the ratio. An empty list is created for each *ei*. When the ratio *n2/n1* for an (*ei*, *ej*) pair is larger than *v2*, we add *ej* to the corresponding list for *ei* and record the lag in the CCG bin with the largest number of counts as the delay time of *ej* relative to *ei*. After the list is constructed, for each *ei*, if all *ej* in the list have non-negative delay times, we store this list as a set of candidate electrodes for a propagation. The requirement for non-negative delay times for all *ej* avoids duplicate detection of the same eAP propagation. This process is repeated for all electrodes *ei* in an array. In this paper, we chose *v2* to be 0.5. The value 0.5 was determined empirically with results from different thresholds compared with manually detected propagation signals for our dataset as described in Tovar et al., 2018. Users can change *v2* to any number between 0 to 1 based on how sharp they want the peaks for a qualified electrode pair to be.

After a set of candidate electrodes is found for a reference electrode *ei*, the second step is as follows. First, find the electrode *eh* in the set of candidate electrodes with the maximum number of co-occurrences with *ei*. If there are multiple electrodes with the same number of co-occurrences with *ei*, the one with a shorter delay time is identified as *eh*. If there are multiple electrodes with the same number of occurrences with *ei* and the same delay time, the one with a smaller ID number is used as *eh*. Second, scan through all other electrodes in the set of candidate electrodes to identify only the electrodes with more than *v3*% of the maximum number of co-occurrences and an *n1* greater than *v4* (*n1* denotes the largest sum of counts in any 0.5 ms moving window in the CCG). Action potential propagation is a high-fidelity process where ideally, all cohort electrodes detect most of the propagating eAPs. In practice, missed spike detection happens in electrodes with low signal-to-noise ratio. *v3* is used as the threshold for the lower bound of the percentage of co-occurrences each cohort electrode needs to detect for an eAP propagation. Given the low inter-electrode latency variability, *n1* is a proxy of the absolute number of co-occurrences. Co-occurrences can happen from random alignment. However, we showed in Figure 2G and 2H that random alignment only leads to a small number of co-occurrences. The threshold *v4* sets a lower bound for the number of co-occurrences to avoid falsely identifying an electrode as a cohort electrode as a result of random alignment.

In this paper, we used *v3* = 50 and *v4* = 50. *v3* was determined empirically with results from different values compared with manually detected propagation signals for our datasets as described in Tovar et al., 2018. We chose *v4* to be 50 based on the number of spikes in each recording. Electrodes that satisfy these criteria along with *ei*, *eh* are identified as the cohort electrodes of an eAP propagation. This process was repeated for all sets of candidate electrodes. This generates a collection of cohort electrodes, each representing an eAP propagation in each recording.

All *vi* can be defined by users.

### Signal statistics extraction

The eAPs from the identified cohort electrodes can be used to extract single cell statistics. Spike times *Ti* for an identified neuronal source *si* are computed using the spike times on the cohort electrodes. Let *ei* again denote the reference electrode in the cohort and *eh* denote the electrode with the most co-occurrences with *ei*. Here co-occurrences were spikes at cohort electrodes that occurred within 1.5 ms window following the eAP at the reference electrode. *ei* and *eh* are the first two anchor points for *si*. First, all the co-occurrences between these two anchor points are identified and added to a spike set *S*. Additional anchor points can be added in the order of most to least co-occurrences with anchor point 1. The co-occurrences between the additional anchor point with anchor point 1 are added to *S* to comprise an amended set of propagating eAPs. Note that because inter-electrode latencies are usually small (<1 ms), we use the spike times at anchor point 1 to represent the spike times of the neuronal sources.

In this work, the propagation eAPs were isolated with three anchor points when possible, unless otherwise specified. In cases with only two cohort electrodes, the propagation eAPs were isolated with two anchor points.

The analysis for computational time in Figure 5C was done on a server with the following specifications: Chassis: Gigabyte R282-Z91-00 Rack Mount Server; Motherboard: Gigabyte MZ92-FS0-00; CPUs: two AMD EPYC 2 Rome 7502 32 Core CPUs; RAM: 1TB DDR4 Reg ECC RAM; Operating System: Ubuntu 20.04 LTS. The software used to run the script was MATLAB R2018b. The code used parfor (parallel) for-loops and employed 12 workers on the server for parallel computing.

### Primary cell culture

All animal protocols and procedures described in this study were approved by the Institutional Animal Care and Use Committee (IACUC) of University of California, Santa Barbara and were performed in accordance with the NIH Health Guide for the Care and the Use of Laboratory Animals. All animal experiments were performed in accordance with ARRIVE guidelines. Primary hippocampal neurons were isolated from postnatal day 0 (P0) C57BL/6 mice using previously described protocols. Up to 3 mouse pups were used per round of cell culture and the neurons were pooled for plating on multiple MEAs. Cleaned and UV sterilized MEAs (120MEA100/30iR-ITO arrays; Multi Channel Systems) were coated with 0.1 mg/ml poly-L-lysine (Sigma-Aldrich) for 1 h at 37 °C, rinsed 3 times with sterile water and air dried prior to plating. All recordings were done from cultures that were plated twice. In the first plating, primary cells were plated and allowed to grow and proliferate. After at least a week, another round of primary cells was plated. Cells were plated at 100,000 to 125,000 cells for the first plating and 125,000 to 200,000 for the second plating. For cultures grown on CMOS arrays, cultures of primary glial cells were maintained in separate T-75 flasks, were plated at 150,000 cells per well on MEAs and allowed to proliferate for at least a week. Freshly dissociated hippocampal cells were then plated at 250,000 cells per dish (550 cells/mm^2^) on the confluent glial cultures. All primary mouse cultures were grown in minimum essential medium with Earle’s salts (Thermo Scientific, catalog # 11090081) with 2 mM Glutamax (Thermo Scientific), 5% heat-inactivated fetal bovine serum (Thermo Scientific), and 1 ml/l Mito + serum extender (Corning) and supplemented with glucose to an added concentration of 21 mM.

### Human brain cerebral organoid generation

Details on generation of the cerebral brain organoids have been previously published (Sharf et al., 2021). The control induced pluripotent stem cell (iPSC) line F12442.453 were cultured in mTeSR1 medium (Stem Cell Technologies) using hESC-qualified Matrigel-coated tissue culture plates (Corning). Media was exchanged every other day and iPSCs were routinely passed using ReLeSR (Stem Cell Technologies).

Organoids were generated using the method of Lancaster et al. (2013), with minor modifications. For dissociation, iPSCs were incubated in 0.5 mM EDTA in D-PBS for 3 min followed by Accutase for 3 min at 37 °C and triturated to achieve a single-cell suspension. Cells were centrifuged for 3 minutes at 1,000 rpm, resuspended in low bFGF hES media supplemented with Rock inhibitor (50 μM) and plated in U-bottom ultra-low attachment plates at 4,500 cells per well. On day 2, low bFGS hES media with Rock inhibitor was replaced. Media was changed on day 4, omitting bFGF and Rock inhibitor. Embryoid bodies were transferred to neural induction media on day 5 (1x N2 supplement, 1x GlutaMAX, 1x MEM-NEAA, 1ug/ml Heparin in DMEM/F12) and media was replaced on days 7 and 9. On day 10, each neuroepithelial structure was embedded in 15 ul hESC-qualified Matrigel, followed by a 2 day incubation in neural induction media.

On day 12, neural induction media was replaced with NeuroDMEM-A media (0.5x N2 supplement, 1x B27supplement without Vitamin A, 1x β-Mercaptoethanol, 1x GlutaMAX, 0.5x MEM-NEAA, 250 ul/l insulin solution, 1x Pen/Strep in 50% DMEM/F12 and 50% Neurobasal). After day 19, organoids were cultured in NeuroDMEM+A media (0.5x N2 supplement, 1x B27 supplement with Vitamin A, 1x β- Mercaptoethanol, 1x GlutaMAX, 0.5x MEM-NEAA, 250 ul/l insulin solution, 12.5 mM HEPES, 0.4 mM Vitamin C, 1x Pen/Strep in 50% DMEM/F12 and 50% Neurobasal) with media changes twice per week. After day 21, organoid cultures were kept on an orbital shaker at 75 rpm.

### Recording conditions

All recordings from low density arrays (MultiChannel Systems; Reutlinger, Germany) or CMOS arrays (Maxwell Biosystems; Zurich, Switzerland) were done in cell culture medium to maintain sterility (see above). Low density arrays (100 micron inter-electrode distance) were recorded using MultiChannel Systems MEA 2100 acquisition system. Data were acquired at 20 kHz and post-acquisition bandpass filtered between 200 and 4000 Hz. Data were acquired on all 120 data channels. We controlled the head stage temperature with an external temperature controller (MultiChannel Systems TC01). Most recordings reported here were done at 30° C, unless otherwise indicated. The media osmolality was usually ∼320 mosmol. Salts were obtained from Sigma-Aldrich or Fluka. Any drugs, such as TTX (Tocris), were introduced into the recording chamber in a volume not more than 0.2% of the chamber volume.

All recordings were done on neurons at 5-30 days in vitro (DIV) except for recordings done on organoids. We used recordings with signals present on the majority of channels. Recording duration was kept brief (3 to 5 minutes) to minimize to avoid large changes in CO_2 and pH_ (Tovar et al., 2018) and to minimize data file size. All recordings were done with the MEAs chambers covered by a CO2-permeable, water vapor- impermeable membrane (Potter and DeMarse, 2001) to minimize evaporation, maintain cell culture sterility and decrease media degassing. Membranes were held in place over the recording chamber by a Teflon collar placed over the culture chamber. Once placed in the headstage, each array was allowed to equilibrate to head stage temperature for at least 5 minutes. For experiments requiring temperature changes, head stage temperature was monitored and each MEA was kept at the new temperature for at least 5 minutes before data acquisition. For all CMOS recordings, the headstage was kept in a cell culture incubator equilibrated to 5% CO2 and 37 °C. CMOS data were acquired at 20 kHz. Data were acquired on a subset (1024 electrodes) of the 26,400 available electrodes.

Recordings from cerebral organoids were performed using a high-density Neuropixels CMOS shank (Neuropixels, Heverlee, Belgium) on 6-month-old cerebral organoids. Briefly, organoids were cultured in BrainPhys media for 30 days prior to recording. For recordings, organoids in BrainPhys media were immobilized in a custom well and kept at 37 °C using a temperature-controlled stage. A 10 mm long high-density Neuropixels CMOS shank (with 384 addressable electrodes in 960 electrode array) was inserted into the organoid using a motorized micromanipulator (MP-285, Sutter Instruments). Data acquisition was done with SpikeGLX (Bill Karsh, https://github.com/billkarsh/SpikeGLX) at a sampling rate of 30 kHz. Subsequent data processing and analysis were performed similarly to 2D CMOS data.

### Spike Detection and Analysis

MultiChannel Systems proprietary files were converted to HDF5 file format prior to all analysis with the MCS program Multi Channel DataManager. Extracellular voltage records were bandpass filtered using a digital 2nd order Butterworth filter with cutoff frequencies of 0.2 and 4 kHz. Spikes recorded from the MultiChannel Systems acquisition system were detected with MEA Tools (Bridges et al., 2017) using a threshold of 6 times the standard deviation of the median noise level (Quiroga et al., 2004). For CMOS recordings, raw voltage records were bandpass filtered between 300- 6000 Hz and the spike detection threshold was 5 times the root mean squared of the signal per channel. Raw data was processed using proprietary MaxLab Live software (Maxwell Biosystems). Manually detected propagating eAPs were initially detected by eye with the help of MEA Viewer (Bridges et al., 2017) spike visualization software and validated by signal averaging. No spike sorted data was used in our analysis. Some analysis was done using custom software written in Python and Mathematica (Wolfram) and in Igor (Wavemetrics). For example, to compare the output of our detection algorithm with raw data (Figure 1), output files from MEA Tools containing spike times and amplitudes were manually compared to amplitude histograms from eAPs isolated with our algorithm. All statistical data are displayed as the mean ± standard deviation.

### Data Availability

All data are available upon request from the corresponding author. Data in this work will be uploaded to the Collaborative Research in Computational Neuroscience data sharing website (http://crcns.org/).

### Code Availability

Code for our algorithm is available at https://github.com/ZhuoweiCheng/Propagation-Signal-and-Synaptic-Coupling-Algorithm

## Supporting information

## Acknowledgments

This study was funded by the Arnold O. Beckman Postdoctoral Fellowship Award and Larry L. Hillblom Foundation: 2019-A-013_SUP

