## Supplementary Figure 1 for "Automated detection of extracellular action potentials from single neurons"

**Supplementary Text**

Each panel shows the amplitude distribution of all eAPs at an electrode (in grey) superimposed with the amplitude distribution of propagating eAPs (red) isolated using different thresholding. The first three panels display how thresholding, based on different multiples of the standard deviation (α), affect the isolation of propagating eAPs. The final panel shows the isolated propagating eAPs amplitude distribution without any such thresholding. The amplitude distribution at this example electrode is divided into two modes; we expect the propagating eAPs to be limited to an apparent single mode. However, a small number of spikes from the lower amplitude mode were included by the algorithm. With different value of α, different number of eAPs were removed. Without thresholding, there are 1939 propagating eAPs in the higher amplitude mode and 139 propagating eAPs in the lower amplitude mode. When threshold was set to $\alpha = 2$, 90% of the eAPs in the lower amplitude mode were filtered out while retaining all of the eAPs in the higher amplitude mode. In general, smaller α will filter out some true co-occurrences and cause higher false negative rate whereas a larger α can cause higher false positive rate.

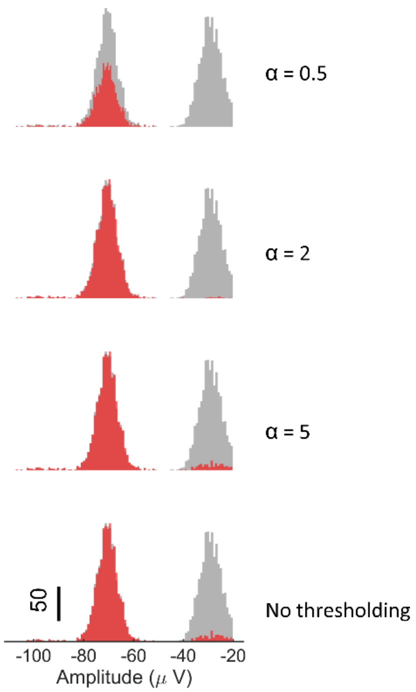


**Supplementary Figure 1.** An example of the usage of multiples of standard deviation (α) of the latency distribution for thresholding. In each histogram, grey are the amplitudes of all eAPs at this electrode. Superimposed on these histograms are the amplitude distributions of isolated propagating eAPs (red). From top to bottom, the first three histograms show the eAP amplitude distribution after thresholding with α = 0.5, α = 2 and α = 5 respectively.  The histogram at the bottom is the propagating eAPs detected without any thresholding.
